# Prioritizing Small Molecule as Candidates for Drug Repositioning using Machine Learning

**DOI:** 10.1101/331975

**Authors:** Khader Shameer, Kipp W. Johnson, Benjamin S. Glicksberg, Rachel Hodos, Ben Readhead, Max S. Tomlinson, Joel T. Dudley

**Affiliations:** Institute of Next Generation Healthcare, Department of Genetics and Genomic Sciences, Icahn School of Medicine at Mount Sinai, Mount Sinai Health System, New York, NY; Current affiliation: Center for Research Informatics and Innovation; Department of Information Services, Northwell Health, New York, NY

## Abstract

Drug repositioning, i.e. identifying new uses for existing drugs and research compounds, is a cost-effective drug discovery strategy that is continuing to grow in popularity. Prioritizing and identifying drugs capable of being repositioned may improve the productivity and success rate of the drug discovery cycle, especially if the drug has already proven to be safe in humans. In previous work, we have shown that drugs that have been successfully repositioned have different chemical properties than those that have not. Hence, there is an opportunity to use machine learning to prioritize drug-like molecules as candidates for future repositioning studies. We have developed a feature engineering and machine learning that leverages data from publicly available drug discovery resources: RepurposeDB and DrugBank. ChemVec is the chemoinformatics-based feature engineering strategy designed to compile molecular features representing the chemical space of all drug molecules in the study. ChemVec was trained through a variety of supervised classification algorithms (Naïve Bayes, Random Forest, Support Vector Machines and an ensemble model combining the three algorithms). Models were created using various combinations of datasets as Connectivity Map based model, DrugBank Approved compounds based model, and DrugBank full set of compounds; of which RandomForest trained using Connectivity Map based data performed the best (AUC=0.674). Briefly, our study represents a novel approach to evaluate a small molecule for drug repositioning opportunity and may further improve discovery of pleiotropic drugs, or those to treat multiple indications.

## INTRODUCTION

Developing a new drug requires significant investments including time, human labor, and resources. A typical drug discovery project takes approximately 15 years with an estimated budget of USD $800- $1 billion [1]. Further, it involves multiple phases including pre-clinical development, which primarily focuses on understanding a biological entity driving an underlying pathology or pathway associated with a disease. Once a candidate target is identified and selected, various medicinal chemistry and bio-pharmacological assessments (i.e. chemical screening) lead small molecule identification, biological experimentation by *in vitro* and *in vivo* studies, bioavailability, dosing, formulation, and extensive multi-center clinical trials testing. In addition to potential failures at every step in the process, often some unintended side effects are found in later phases of development and post-market phases (i.e. side effect due to drug-drug interaction). As multiple factors influence success in drug discovery, novel approaches are required to deliver therapies to market in short turnaround timelines, particularly for disorders in which there is no robust standard of care. Drug repositioning is a unique strategy by which drug discovery researchers and pharmaceutical companies can control the cost by leveraging the library of Food and Drug Administration (FDA) or other regulatory body approved medications for new indications [2]. Systematic drug repositioning evaluates a library of approved or investigational pharmaceutical compounds as a therapeutic modulator for a new disease indication. Repositioning was reported in various classes of disorders defined using International Statistical Classification of Diseases and Related Health Problems (ICD-10). Multiple therapeutic areas have examples of successful drug repositioning. Drug repositioning supported by comparative efficacy trials could deliver a drug to a market faster than traditional drug discovery cycle. To date, more than 250 drug-repositioning investigations have been published in the biomedical literature, suggesting reuse of medications for common, rare or orphan diseases is a viable and pragmatic approach [3–5]. However, the ability to assess whether a drug might be effectively be repositioned to a new indication is not currently addressed and represents a knowledge gap. In this manuscript, we present a novel chemoinformatics based machine learning model to address this need; specifically, to predict the suitability of small molecules as candidates for drug repositioning.

### Previous work

Previously, three-dimensional structure prediction of target proteins, protein-drug based docking scores, and biomedical knowledge-mining using text-mining algorithms were used to predict and prioritize compounds for potential drug repositioning, *de novo* drug discovery, or improving the quality of small molecules [6–8]. Recently, Napolitano et al. showed that predicting the probability of compounds being repositioned could be enhanced by integrating gene expression data, chemical information, and target information [8]. While these methods are useful, they require a significant amount of data types for predictive analytics. Measuring such diverse modalities of compounds in the initial stages of drug discovery is challenging. Machine learning has been applied to various problems in biomedicine and healthcare and can be employed for this problem. Due to the recent advances in machine learning, coupled with growing availability of large datasets and more affordability of computing resources, different learning tasks in drug discovery has been met with varying degree of success. Artificial intelligence methods including machine learning techniques including deep learning has been used to find binding sites and therapeutic conformations of peptides, as well as to prioritize compounds for experimental studies and clinical evaluation [3, 4]. However, no comprehensive machine-learning model is available to evaluate the suitability for drug repositioning opportunities for a given compound using minimal chemical information.

## METHODS

In the following section, we discuss the datasets, feature engineering strategies, and machine learning approaches utilized in the study. We present a brief outline of the methodology used in this study in Figure 1.

**Figure 1:**
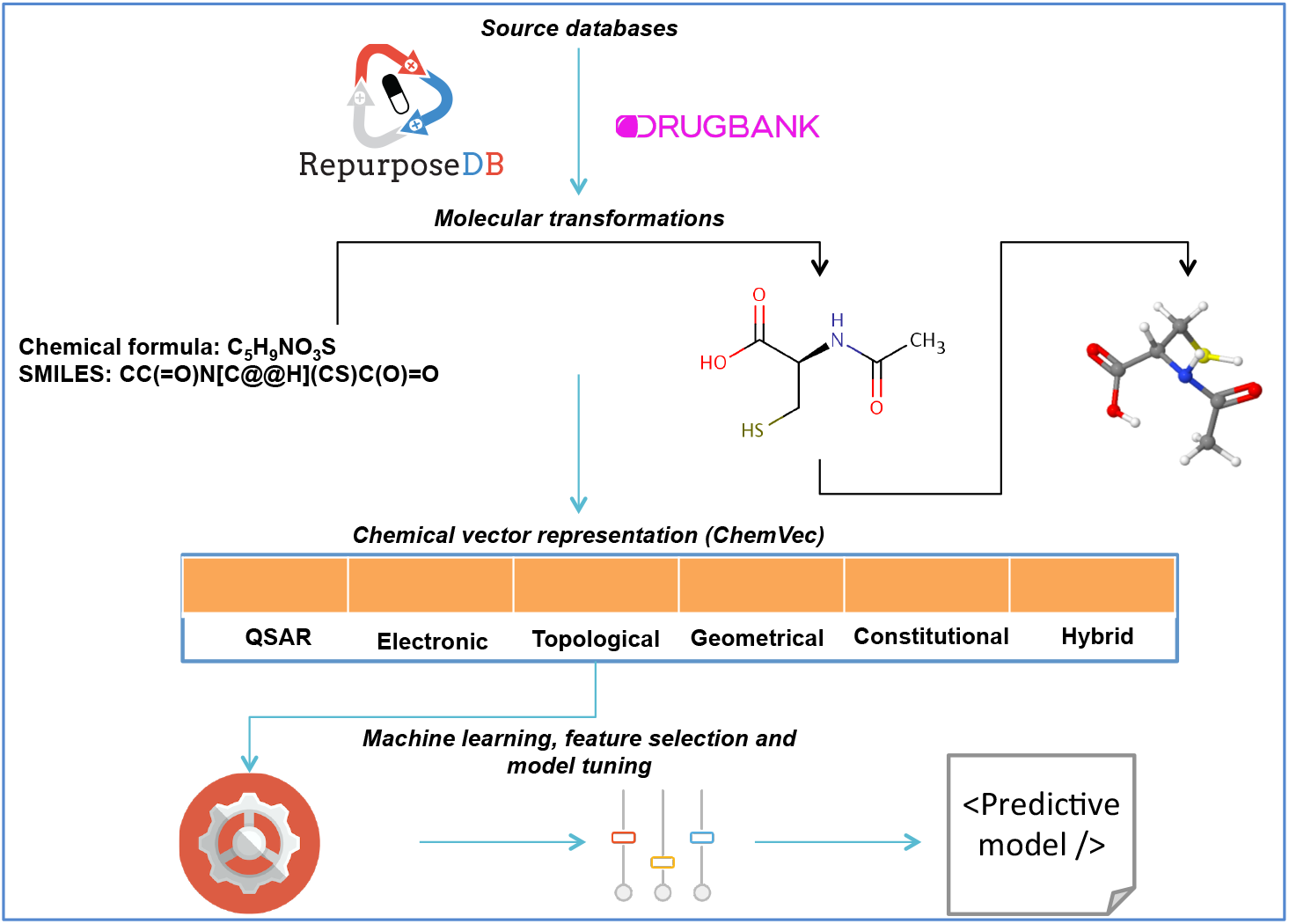
Brief outline of modeling strategy used to predict drugs suitable for repositioning

### Dataset

RepurposeDB (http://repurposedb.dudleylab.org) is a compendium of successful drug repositioning investigations that we previously developed to address a critical knowledge gap in computational drug discovery [9]. The motivation for developing RepurposeDB is to catalog repositioned drugs (small molecules and protein drugs; n=253) with their primary indications and secondary indications. Briefly, RepurposeDB is a multimodal knowledgebase pertaining to repurposed drugs, including their known chemical and biological information. We further explored other data types including shared genetic architecture, phenomic similarities, and digital epidemiology data in the form of primary-secondary indication based comorbidity analyses using electronic medical records. Thus, RepurposeDB represents one of the highly annotated, comprehensive reference database on drugs, disease indications with biochemical, pharmacological and epidemiological data related to drug repositioning. In this study, we compiled the drugs (small molecules) subset of drugs from RepurposeDB (n=188) as the positive dataset (repurposed compounds) and the compounds without evidence for drug repositioning in DrugBank [10] that are not in RepurposeDB as the negative dataset. The approval status of these compounds was also considered as an additional filter to fine tune the model to output approved compounds only instead of the entire chemical space in DrugBank (Approved, *n*=1560; Not approved, *n*=5274). It should be noted that approval status is at the time of this analysis and is subject to change over time. Predictive learning was performed using three different versions of the negative dataset: DrugBank full (DrugBank-F, *n*=6647) approved subset of DrugBank (DrugBank-A; *n*=1374) and subset of compounds in Connectivity Map (DrugBank-C; *n*=631).

### ChemVec - feature engineering using chemical information

Chemical features (e.g. ligand classes) represent several opportunities for feature engineering. For example, a single chemical compound can be represented in multiple modalities as one-dimensional alphanumeric representations (for example: chemical formulae or simplified molecular-input line-entry system [SMILES] representation). Chemical molecules can also be represented as three-dimensional structures in atomic resolution. Further the molecular dynamics nature of drug-drug, drug-protein, and drug-environment interactions can be modeled in a vacuum or an aqueous medium. Each of these representations provides multiple types of data: for example, using molecular formulae of SMILES representation we can compute atomic and molecular composition. SMILES encode connectivity information but not using three-dimensional structure features like accessible surface area. Coupling three-dimensional structure features and molecular dynamics information would help to compute various geometric parameters like kier shapes and Zagreb group indices. We compiled chemical descriptors of the small molecules (i.e. quantitative structure activity relationship parameters, electronic, topological, geometrical, constitutional and hybrid) using the following libraries: chemistry development kit Pybel [11] and JOELib, and combined using Python and R scripts. We call the unique chemical signature framework derived from the combination of features from different molecular representations as a chemical information vector space: “ChemVec”. For all predictive models, the ChemVec molecular space comprising 111 features is ranked and ChemVec features contributing to the prediction tasks are elucidated. We compiled and tabulated the data, as well as performed all machine learning tasks, using R package *caret*. We derived feature importance using information gain criterion and ranked using Ranker method, implemented using the Weka workbench. For each chemical structure, we converted the chemical formula from the SMILES format into a 2D and 3D model to generate structure files. We then optimized the 3D models using 500 iterations of local energy minimizations with MMFF94 force field to represent a native confirmation of the chemical structures.

### Machine learning – parameterization and evaluation of algorithms

We trained the following machine learning models to predict drug repositioning from chemical information: Naïve Bayes (NB), Support Vector Machines (SVM), RandomForests (RF), and Ensemble Model (EM) combining three different algorithms (NB, SVM and RF with majority voting). We have implemented NB, SVM, RF and EM as explained in our previous studies using standard parameterization suitable for biomedical datasets and applied using R [12, 13]. Models were tested using 70% of the dataset and trained using 30% of the dataset. All predictions were evaluated using parameters including precision, recall and accuracy rates.

## RESULTS

Feature selection using Information Gain methods suggests the following features relevant for different predictive models: Connectivity Map based model: LogP, C, Geometrical diameter, Molecular Refractivity, Kier shape 2, Number of bonds, Number of atoms, AROMATIC, RINGS, abonds, Geometrical shape coefficient, Number of heavy bonds, Number of P atoms, Number of HBD 1, Number of HBD 2, Number of HBA 2and Pt; Approved Compounds based model: LogP, C, Kier shape 1, bonds, RCOR, AROMATIC and Fe; DrugBank Full based model: RCOR, Fraction of rotatable bonds, ROPO3, ROR, Number of acidic groups, AROMATIC, RCHO, P, RCN, As, Al, tbonds and Pt Evaluation of a total of 15 models using three negative datasets reveals RandomForests with DrugBank-C as the model with highest accuracy rate (Table 1). We also noted that Connectivity Map based model had the lowest number of compounds yet the highest number of features contributing to the prediction task. Three features (LogP, C, and bonds) were predictive across three models and they were independently validated in our previous work as chemical features driving drug repositioning and hence provide additional orthogonal evidence for the predictive model.

**Table 1:** Summary of predictive models developed and various model evaluation metrics

| Models | Dataset | Algorithm | Precision | Recall | ROC Area |
| --- | --- | --- | --- | --- | --- |
| Connectivity Map | RepurposeDB ($n=187$)<br>DrugBank-C ($n=631$) | Naïve Bayes | 0.632 | 0.698 | 0.660 |
|  |  | SVM | 0.549 | 0.549 | 0.489 |
|  |  | **Random Forest** | **0.735** | **0.763** | **0.674** |
|  |  | Ensemble Model | 0.725 | 0.755 | 0.550 |
| Approved Compounds | RepurposeDB ($n=187$)<br>DrugBank-A ($n=1374$) | Naïve Bayes | 0.806 | 0.312 | 0.519 |
|  |  | SVM | 0.744 | 0.857 | 0.496 |
|  |  | **Random Forest** | **0.780** | **0.859** | **0.612** |
|  |  | Ensemble Model | 0.744 | 0.857 | 0.496 |
| DrugBank Full | RepurposeDB ($n=187$)<br>DrugBank-F ($n=6647$) | Naïve Bayes | 0.956 | 0.903 | 0.582 |
|  |  | SVM | 0.955 | 0.976 | 0.500 |
|  |  | **Random Forest** | **0.955** | **0.976** | **0.597** |
|  |  | Ensemble Model | 0.977 | 0.955 | 0.500 |

## CONCLUSIONS AND FUTURE WORK

In this report, we present the design and development of a predictive model for a challenging task with minimal information from chemical moieties: identifying drugs, which have potential to be repurposed but have not yet been. Drug discovery is a costly endeavor that often takes decades of pre-clinical research, clinical trials, and large cost with the possibility of failure at every step. There are no algorithms or predictive models currently available to test if a compound is suitable for drug repositioning using only chemical information. Our applied machine learning model will allow drug discovery investigators to quickly query ranked and prioritized compounds that could be a candidate for repositioning for any disease indication. We developed ChemVec, a small-molecule-based vector tool as part of this study, which could be beneficial for a variety of chemoinformatics and bioinformatics prediction tasks including molecular interactions, target prioritization, side effect profiling and personalized prescription. We plan to extend ChemVec and build more complex feature representations using deep learning for faster learning. Such models could be trained and tested using massive compound datasets including PubChem to enhance discovery novel compounds capable of drug repositioning opportunities.

